## Supplementary figures and images for "Glycogen phase separation drives macromolecular rearrangement and asymmetric division in *Escherichia coli*"

### Fig EV1

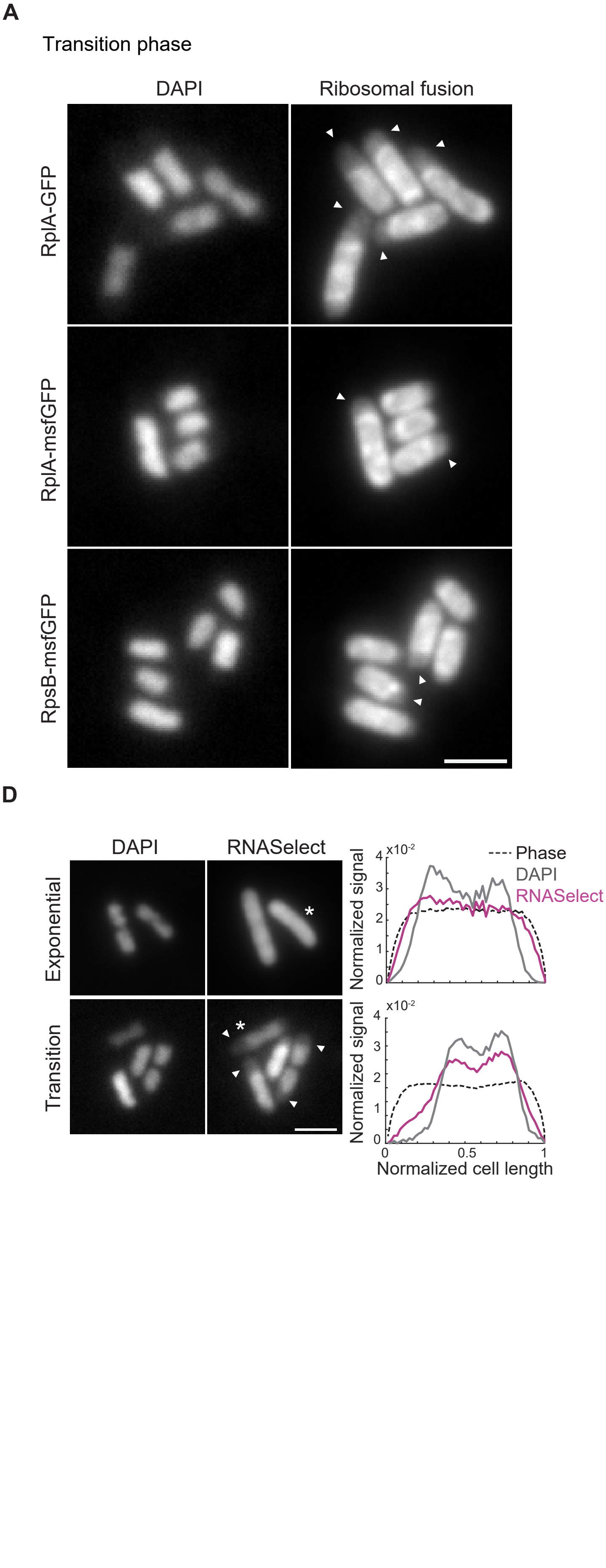

### Fig EV2

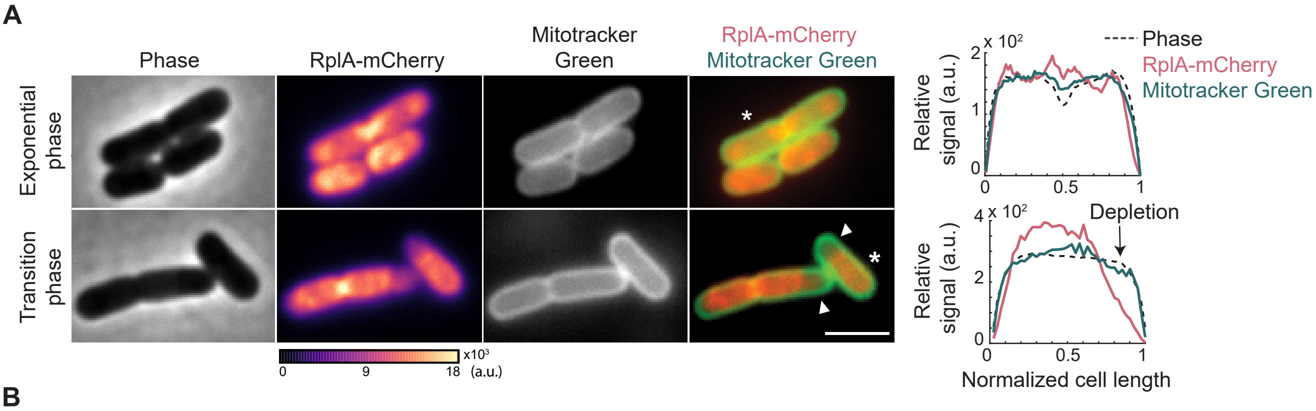

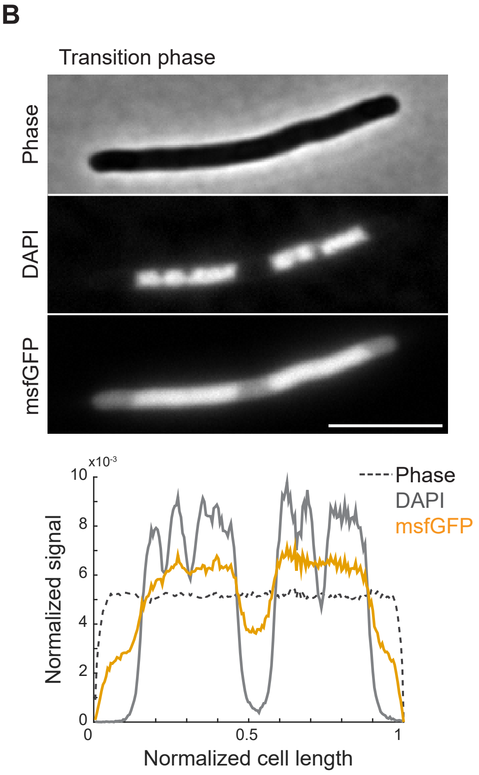

### Fig EV3

Transition phase

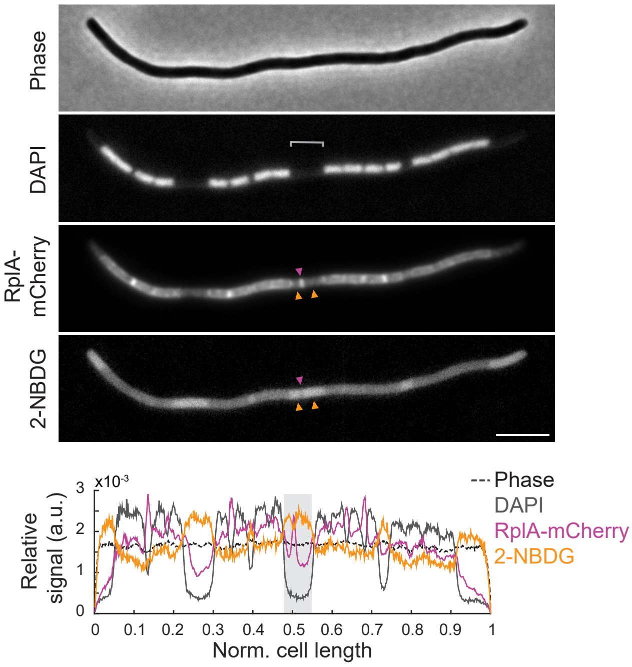


#### Fig EV3

### Fig EV4

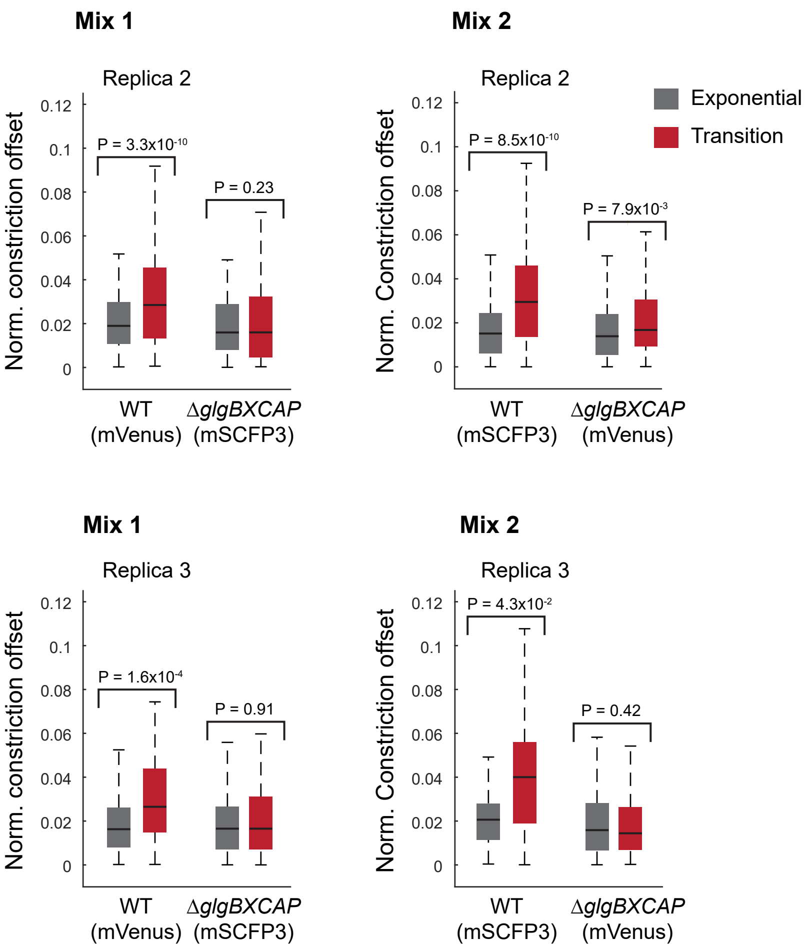

### Fig EV5

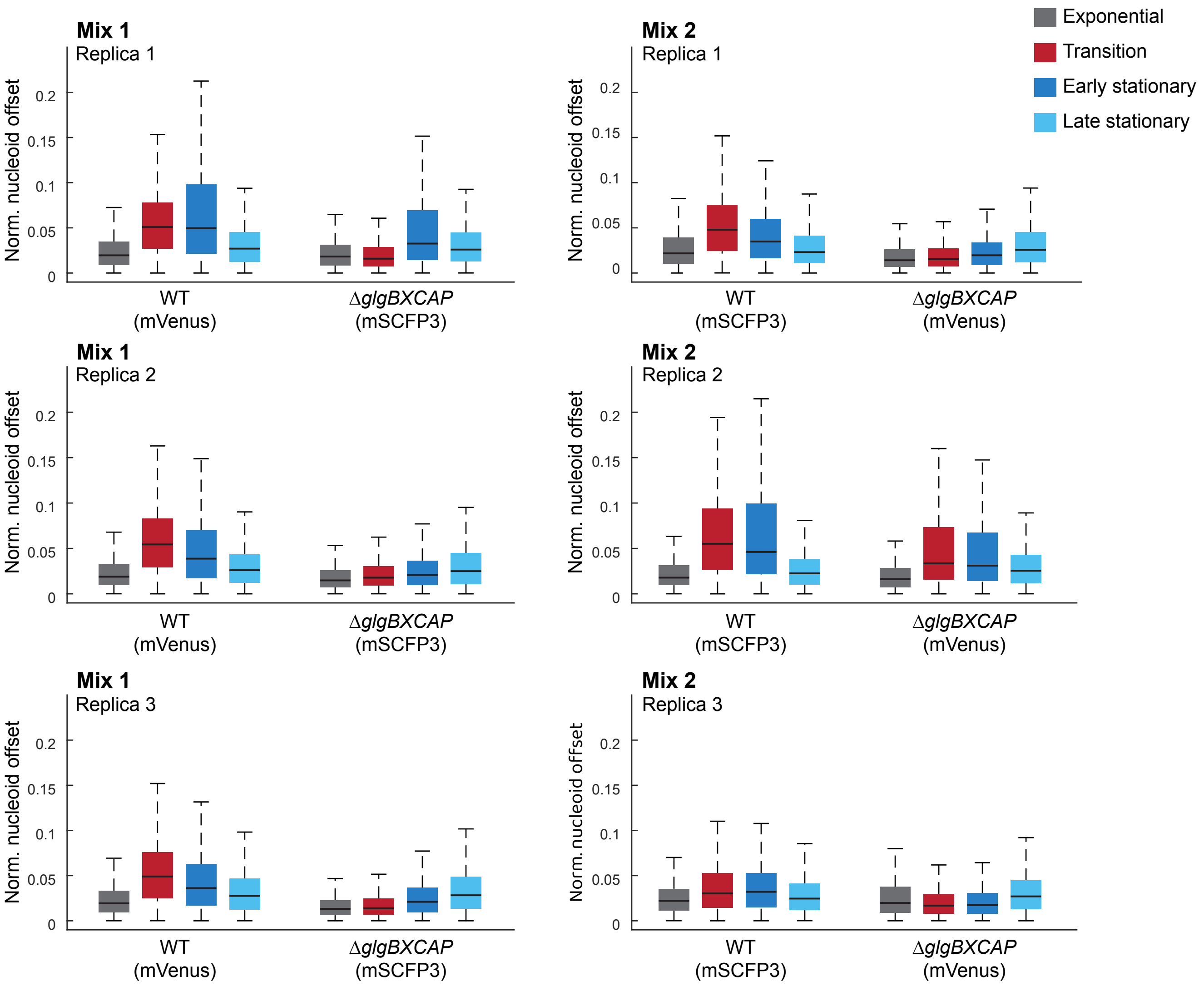

### Fig EV6

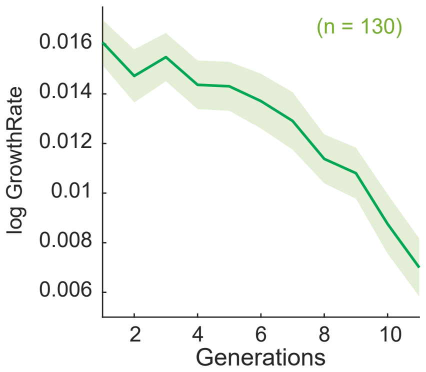

### Fig EV7

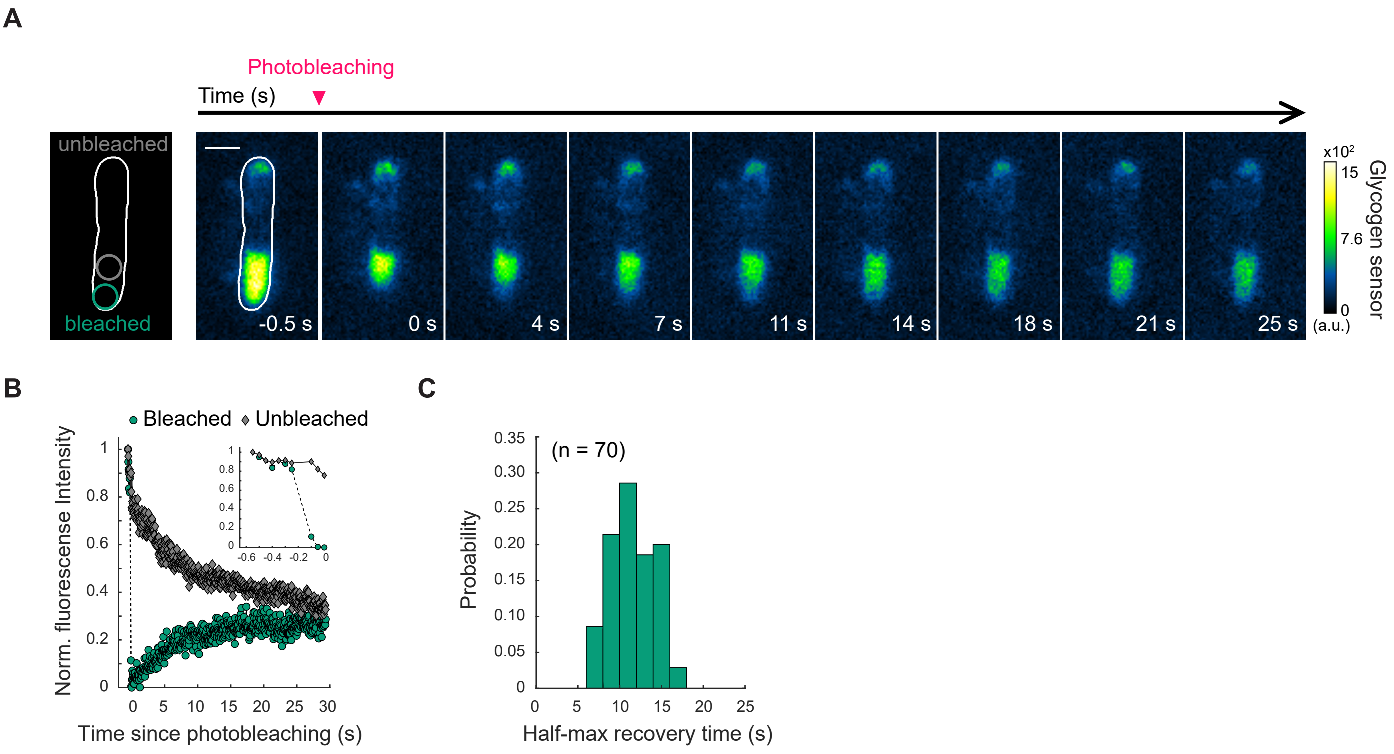

### Fig EV8

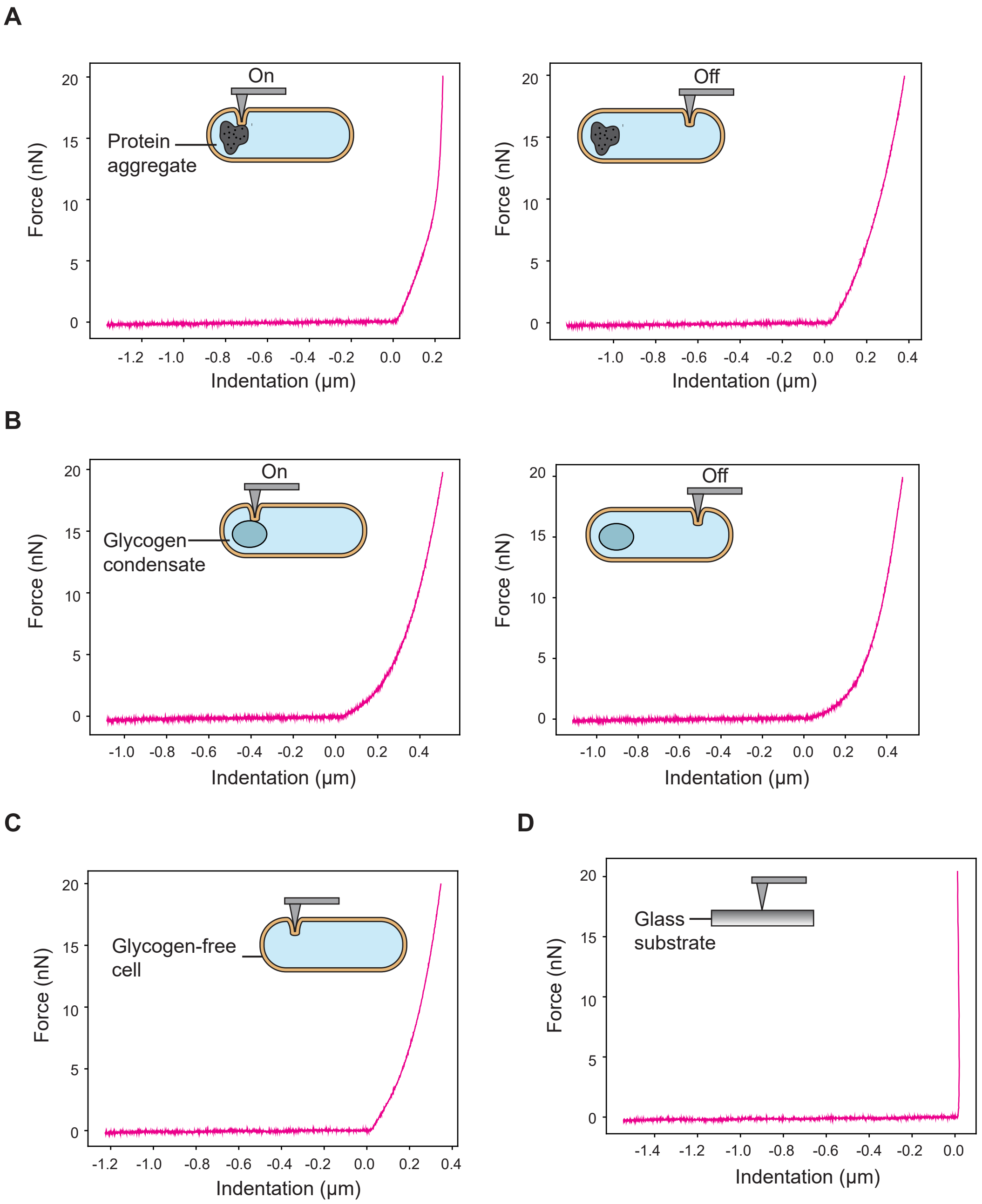

### Fig EV9

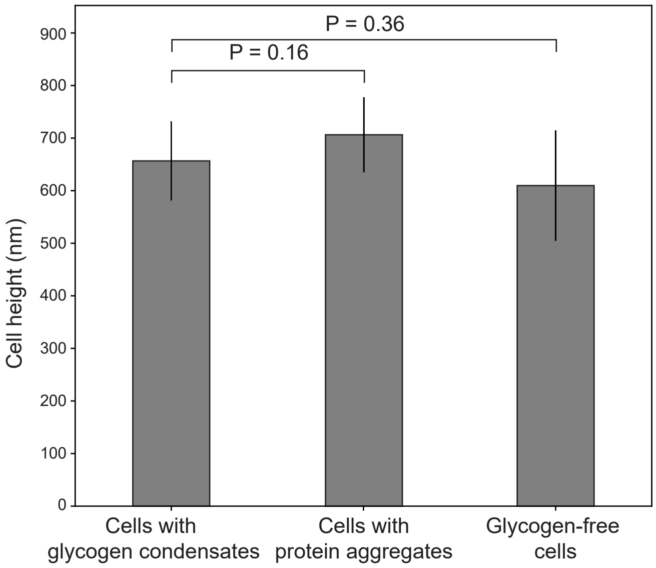

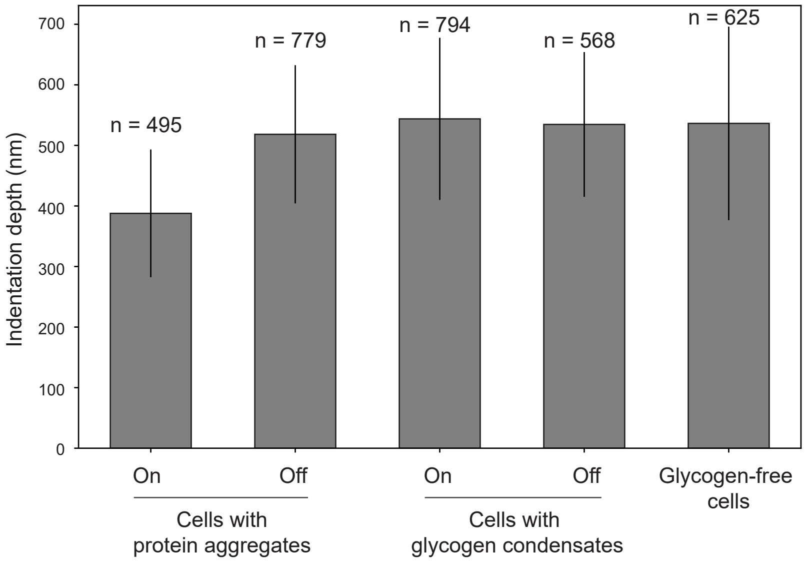

### Fig EV10

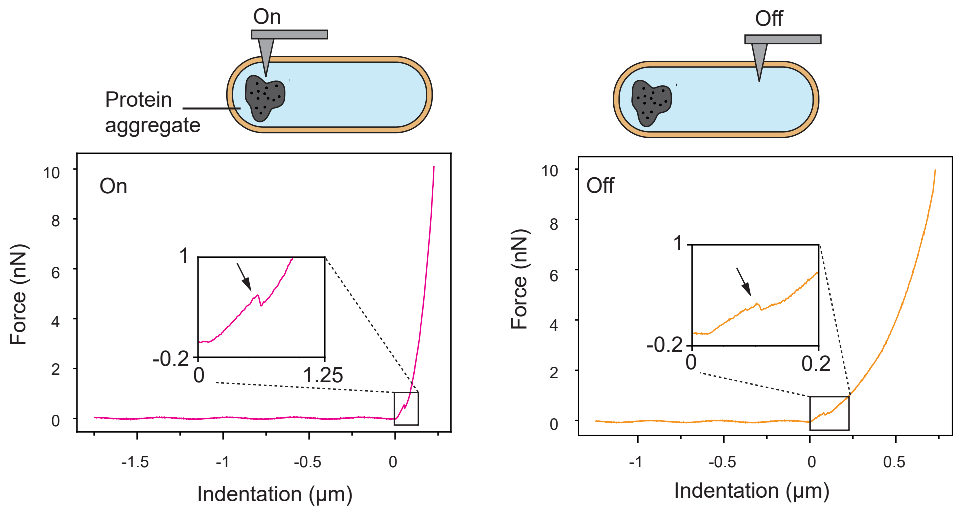

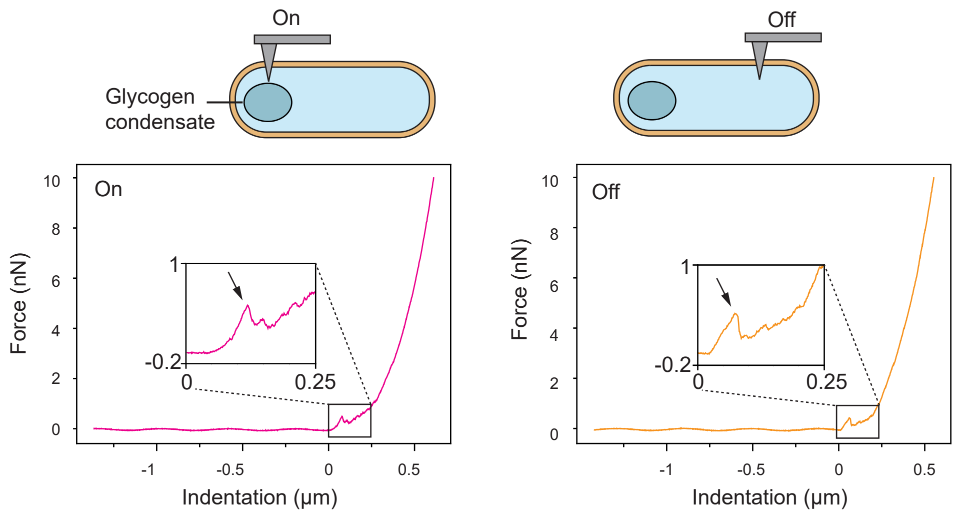

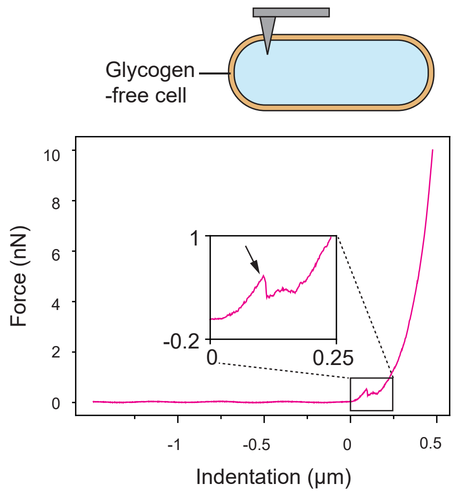
